# PromptBio: An Agentic Platform for End-to-End Computational Biomedical Research

**DOI:** 10.64898/2026.09.02.748774

**Authors:** Minzhe Zhang, Wenhao Gu, Bowei Han, Wenbin Guo, Jiayu Chen, Xiaoyuan Zhou, Yang Leng, Youjia Ma, Kai Li, Junbin Zheng, Hebbar Shishir, Weiying Wang, Aaron Huang, KC Shashidhar, Xiao Yang

**Affiliations:** PromptBio Inc., 7068 Koll Center Pkwy, Suite 402, Pleasanton, CA 94566

**Author notes:** Corresponding authors: KC Shashidhar, Xiao Yang.

## Abstract

Modern biomedical research increasingly depends on complex computational analyses, yet translating a scientific question into a reliable workflow still requires substantial technical expertise and manual coordination. PromptBio is a multi-agent AI platform available through a web portal at https://promptbio.ai that addresses this challenge. Through natural-language interaction, its agent harness, PromptGenie, translates a research objective into an inspectable plan, executes the required research and analysis, and adapts subsequent steps as evidence and results emerge. PromptBio integrates reasoning with managed execution while preserving human oversight and a traceable record of the research process. It can apply validated methods, construct custom analyses, and incorporate external workflows, enabling researchers to move beyond fixed pipelines while supporting reproducibility. We evaluate the platform through benchmarks of end-to-end bioinformatics analysis and biomedical deep research, validation of representative omics and machine-learning skills, and a hypothesis-driven regulatory-genomics case study. PromptGenie achieved higher analytical accuracy and stronger evidence retrieval and synthesis than the comparison agents, while the evaluated domain skills produced results consistent with established methods. The case study further demonstrates how PromptBio can integrate human-specific genomic and epigenomic features to investigate cortical development. Together, these findings show that PromptBio can coordinate complex biomedical analyses under researcher oversight, offering an accessible approach for accelerating scientific iteration and transforming research questions into transparent, reusable computational workflows.

## 1 Introduction

Computational biomedical research requires researchers to coordinate heterogeneous data, specialized analytical methods, and a rapidly expanding scientific literature. A single research question may involve finding relevant studies and public datasets, evaluating evidence, processing multi-omics data, selecting statistical or machine-learning methods, executing interdependent analyses, and interpreting the results in biological context. These activities are commonly distributed across disconnected repositories, software tools, and computing environments. The resulting fragmentation slows scientific iteration, complicates reproducibility, and creates barriers to translating data and literature into coherent, evidence-supported findings.

Large language model (LLM)-based scientific agents increasingly support literature retrieval, hypothesis development, data exploration, workflow generation, and computational or experimental execution [1]. Established cloud bioinformatics platforms such as DNAnexus [2] and Seqera [3] remain centered on scalable data management and workflow execution, while increasingly adding AI-enabled and, in some cases, agentic interfaces to those environments. General-purpose biomedical agents such as Biomni [4], BioMedAgent [5], STELLA [6], and OpenBio [7] plan and execute diverse research tasks through tool use and code generation. Scientific discovery systems such as FutureHouse’s Robin [8] and Google’s Co-Scientist [9] emphasize hypothesis generation and experimental design, while complementary systems such as Noah.bio [10] focus on literature- and database-backed evidence synthesis. These systems demonstrate the potential of natural-language interfaces and agentic methods in biomedical research. An unresolved challenge is to connect evidence retrieval, adaptive planning, executable analysis, and interpretation within a coherent research process while preserving provenance, reproducibility, and expert oversight [11].

PromptBio addresses this challenge through an agentic platform for computational biomedical research. At its center is PromptGenie, an agent harness that interprets a research objective, constructs an inspectable multi-step plan, coordinates specialized subagents and reusable skills, and executes analyses in managed computational environments. Depending on the task, it can combine literature and knowledge retrieval, public or private data access, validated omics pipelines, statistical and machine-learning methods, external workflows, and dynamically generated code. Intermediate evidence and computational results are returned to the harness so that subsequent steps can be revised when inputs are missing, execution fails, or the emerging findings require a different analytical direction.

The platform is designed for human–AI collaboration rather than unrestricted autonomy. Researchers can refine objectives, inspect plans, review analytical choices and intermediate results, and intervene at consequential decision points. PromptBio uses validated tools and domain-standard practices when available; for generated or user-provided analyses, it retains code, environment information, parameters, outputs, and logs to support inspection and reuse. Modular interfaces support extensibility and scalable execution [1], while provenance, monitoring, access control, and compliance-oriented safeguards provide operational support for research involving sensitive biomedical data [12, 13, 14, 15]. These mechanisms support, but do not by themselves guarantee, scientific validity or reproducibility.

This paper presents the design and implementation of PromptBio and evaluates it at complementary levels. We describe the PromptGenie control loop, its coordination of subagents, skills, and execution environments, and the biomedical capabilities available for omics processing, machine learning, knowledge retrieval, and custom workflow generation. We evaluate end-to-end bioinformatics execution using PromptBio-Bench [16], comparing PromptGenie with Biomni and STELLA, and evidence retrieval and synthesis using the Health subset of DeepResearch Bench II [17], comparing PromptGenie with Biomni and Noah.bio. We separately validate representative omics and machine-learning skills against established methods and demonstrate the platform in a researcher-supervised regulatory-genomics analysis of Human Ancestor Quickly Evolved Regions (HAQERs) during cortical development [18]. Together, these analyses assess both the utility of coordinated agentic execution and its current limitations; they do not establish uniform performance across all biomedical domains or fully autonomous research settings.

## 2 PromptGenie: An Agent Harness for Computational Biomedical Research

PromptGenie is the agent harness at the center of PromptBio (Figure 1). Rather than a collection of independent assistants or a router among predefined agents, a harness is an iterative control loop that keeps a research objective in view while gathering context, constructing a plan, delegating specialized work, invoking skills and tools, executing computational steps, evaluating intermediate results, and synthesizing the final response. The same loop can support tasks ranging from focused literature or data queries to long-running, multi-step biomedical research workflows.

**Figure 1:**
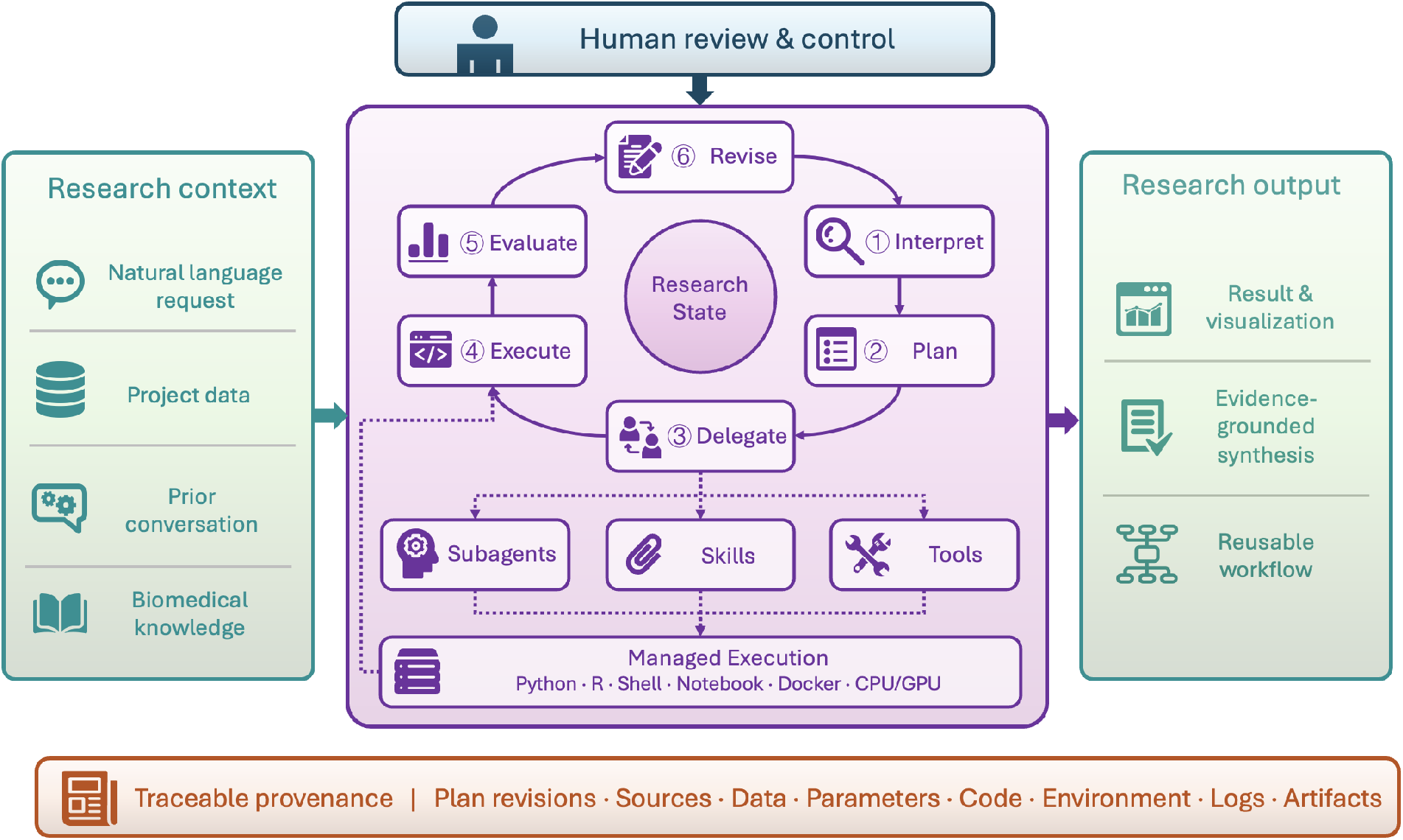
The PromptGenie agent harness. Research context enters an iterative control loop that interprets the request, plans the work, delegates specialized tasks, executes analyses, evaluates intermediate results, and revises the plan as needed. Subagents, reusable skills, tools, and managed computational environments support execution; human review and traceable provenance span the full process, which produces results, evidence-grounded synthesis, and reusable workflows.

PromptGenie maintains the research objective and evolving task state throughout execution. Reasoning and planning provide the control layer; subagents contribute specialized expertise; skills expose reusable research capabilities; and managed execution environments run the required software and workflows. These components operate within a human-in-the-loop process in which users can inspect plans, review intermediate findings, and guide consequential decisions. The resulting architecture combines the flexibility of LLM-based interaction with structured execution, domain-standard methods, and traceable research artifacts.

### 2.1 Reasoning and Planning

Each task begins with PromptGenie interpreting the user’s research objective in the context of available data, prior conversation, project artifacts, and relevant biomedical knowledge. For focused requests, the harness can proceed directly to an appropriate skill or tool. For complex requests, it decomposes the objective into a sequence of research and computational steps, identifies dependencies, and constructs an execution plan. A plan may combine literature retrieval, public-data discovery, data processing, statistical modeling, machine learning, visualization, and biological interpretation within a single task. Information gathering and computation can be interleaved so that newly identified evidence and intermediate results inform subsequent steps and the final synthesis.

Planning is iterative rather than limited to a one-time translation of a prompt. PromptGenie evaluates intermediate outputs and execution status to determine whether the current plan remains appropriate. New evidence, missing inputs, incompatible data, or failed execution can trigger revision of subsequent steps. Dependency-aware plans also allow independent steps to run in parallel while preserving the ordering required for downstream analyses. Users may review and refine the plan before execution and intervene when scientific judgment or additional context is required.

### 2.2 Subagent Orchestration

PromptGenie delegates specialized work to subagents when a task benefits from focused context, domain-specific instructions, or independent execution. A registry describes the capabilities and interfaces of available subagents, allowing PromptGenie to select them according to the requirements of the plan. Subtasks may be executed sequentially or concurrently, and their outputs are returned to PromptGenie for evaluation and integration into the broader research state.

The harness includes subagents for several recurring functional roles. OpsGenie manages projects, users, samples, subjects, datasets, and task bindings while enforcing transactional consistency, auditability, and role-based access control. DataGenie discovers and acquires public biomedical datasets through natural-language and metadata-aware search, including semantic retrieval, structured filtering, and direct cloud-to-platform transfer. It currently indexes fifteen commonly used repositories spanning sequencing, array, proteomics, metabolomics, cancer genomics, and pharmacogenomics resources; dataset and sample counts for each source are summarized in Supplementary Note 1 (Table S1). Additional specialized subagents can assist with omics processing and analysis, custom analysis, knowledge retrieval, and other tasks that require dedicated reasoning or context. PromptGenie remains responsible for the overall plan and synthesis, preventing specialized outputs from becoming isolated from the user’s broader research question.

Subagents complement rather than define the platform’s capabilities. A task does not need to follow a fixed sequence of named agents, and many operations can be performed directly through registered skills. This distinction enables PromptGenie to use a lightweight skill for a well-specified operation while reserving subagent delegation for work that requires additional reasoning, research, or iterative decision-making.

### 2.3 Skills and Tools

Skills provide the reusable capability layer of PromptGenie. Each skill encapsulates the instructions and interfaces required for a defined research operation and may invoke one or more underlying tools. Tools are machine-invocable operations designed for specific purposes, such as knowledge retrieval, data analysis, or result synthesis. Registered metadata describing each skill’s purpose and expected inputs and outputs enables PromptGenie to discover and compose skills during planning. Validated biomedical skills and their integration with external workflows are described in Section 3; the skill library is extensible as new research needs, data modalities, and methods emerge.

When a research question falls outside the predefined skill library, PromptGenie can plan a custom analysis and generate Python, R, or shell code using appropriate scientific packages. The generated code is executed through a code-execution tool in a managed environment; users can inspect the proposed method, the code, and the resulting outputs, and the harness can revise a failed step through the same control loop used for skill-based execution. User- or organization-provided scripts and pipelines can be incorporated as a third path. Together, registered skills, on-demand generation, and imported pipelines allow PromptGenie to apply established methods without constraining researchers to fixed workflows.

### 2.4 Execution Environments

PromptGenie executes computational work in managed environments suited to the requirements of each task. Each step runs in an isolated sandbox that limits generated code and pipeline execution to the assigned environment. The platform supports Python, R, shell, Jupyter notebooks, and domain-specific bioinformatics software, with dependencies provisioned through reproducible environment management. Compute-intensive tasks are assigned to dedicated environments, including GPU-enabled environments when accelerated computation is required. Inputs, parameters, software dependencies, outputs, and execution logs are recorded for each task so that analyses can be inspected and rerun.

Multi-step execution is coordinated by a dependency-aware engine. Steps become eligible for execution when their required inputs are available, allowing independent branches of a workflow to run in parallel. Intermediate artifacts are passed between compatible steps through defined interfaces, while invalid dependencies or incompatible data connections are rejected. This execution model supports workflows ranging from a single generated script to large multi-omics processes that combine preprocessing, downstream analysis, machine learning, and interpretation.

Execution status and errors are returned to PromptGenie as part of the control loop. When a step fails, the harness can inspect the error, revise parameters or code, adjust the plan, and submit the updated step for review or re-execution. Task-scoped access controls protect project data and credentials, while centralized logging and monitoring support troubleshooting, resource management, and operational oversight.

### 2.5 Human Oversight and Provenance

PromptBio is designed to augment rather than replace scientific judgment. Users can review plans, approve or modify analytical choices, inspect generated code, evaluate intermediate results, and redirect the task as evidence develops. Human review is particularly important when the research question is underspecified, datasets or metadata are incomplete, evidence is conflicting, or an analytical decision could materially affect the scientific conclusion. Users also remain responsible for confirming source reliability, determining whether results warrant experimental validation, and ensuring compliance with ethical, privacy, and regulatory requirements.

PromptGenie retains the information needed to reconstruct the research process, including the plan and its revisions, evidence sources, data references, parameters, generated code, execution environments, logs, and resulting artifacts. This provenance connects final statements to the literature, data, and computations that support them. By coupling the flexibility of an agent harness with reviewable plans and traceable execution, PromptBio aims to make complex biomedical research workflows more accessible without sacrificing transparency or reproducibility.

## 3 Biomedical Skills and Pipeline Integration

Building on the skill and tool architecture described in Section 2.3, this section presents PromptBio’s biomedical capability layer (Figure 2). The layer comprises built-in skills for omics processing and analysis (OmicsGenie), machine learning (MLGenie), and literature- and knowledge-based interpretation (MarkerGenie), together with external workflows from nf-core community pipelines, organization-provided pipelines registered through Bring Your Own Pipeline (BYOP), and selected biomedical foundation models.

**Figure 2:**
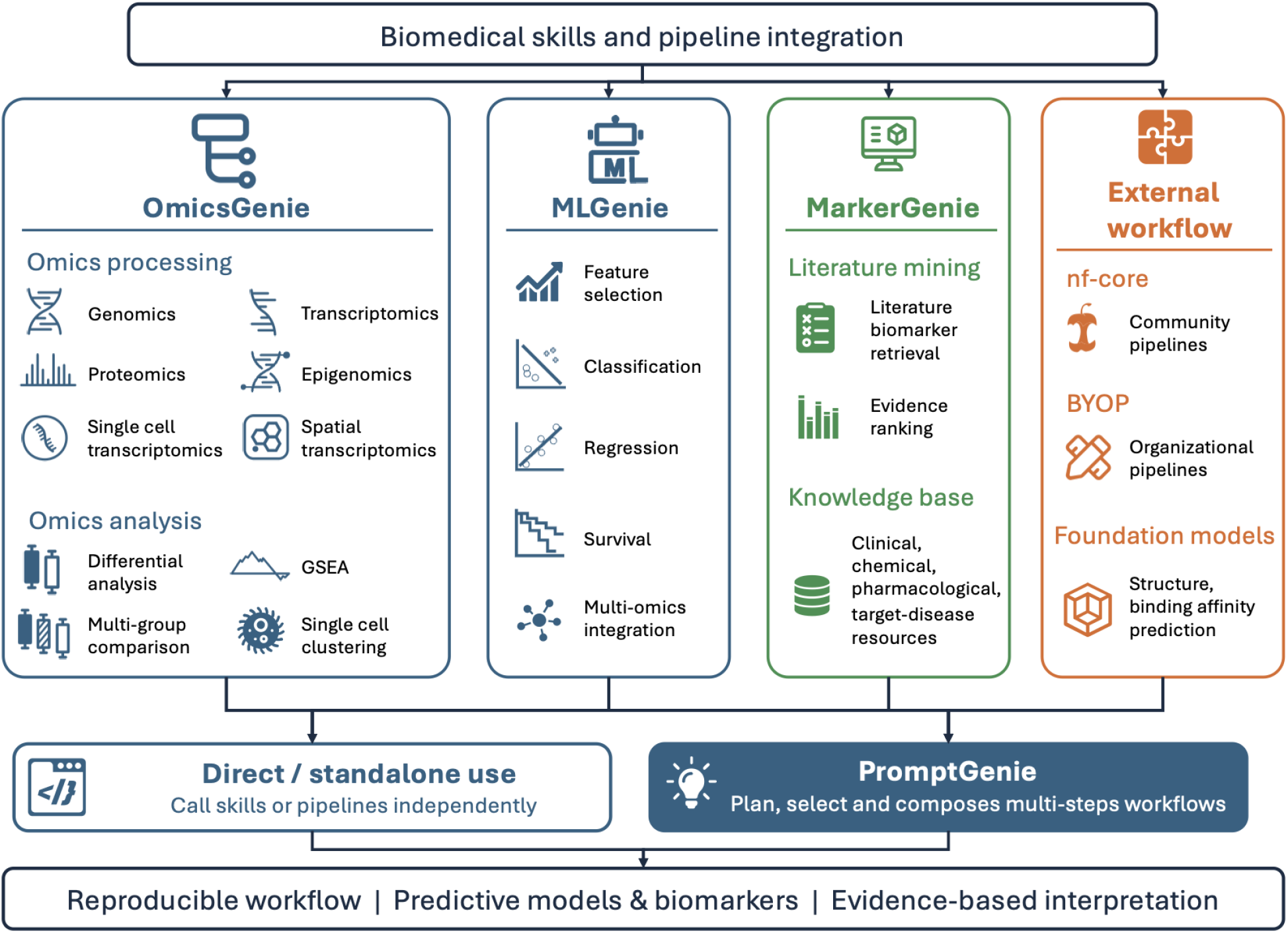
Biomedical skills and pipeline integration. Built-in skills (OmicsGenie, MLGenie, and MarkerGenie) provide validated capabilities for omics processing and analysis, machine learning, and literature- and knowledge-based interpretation. External workflows, including nf-core community pipelines, BYOP, and biomedical foundation models, are integrated through the same capability layer. All resources are exposed through consistent interfaces and can be invoked independently or planned and composed by PromptGenie into multi-step research workflows.

Built-in skills are developed to cover the analyses that arise most frequently in computational biomedical research. They combine established community software with PromptBio-developed modules and are actively maintained so that they remain reliable under agent-driven execution. Each skill exposes a structured representation of the analysis request and of the resulting outputs, making inputs, parameters, and artifacts explicit. Where accelerated computation substantially reduces runtime, execution is further optimized for cost efficiency, including GPU acceleration. These properties make common analyses inspectable to the researcher and allow PromptGenie to select, configure, and compose them rather than reconstructing them for each task.

PromptBio also integrates external resources that are not covered by built-in skills, including community pipelines, user-provided pipelines, and foundation-model services. These resources are described with the metadata required for discovery, configuration, and execution in managed environments, so that they can be reused alongside built-in skills.

Built-in and external capabilities are exposed through consistent interfaces. Each can be invoked independently as a standalone analysis, or planned, selected, and composed by PromptGenie into a multi-step research workflow.

### 3.1 Omics Processing Skills

OmicsGenie provides validated pipelines that transform raw omics data into analysis-ready outputs. The genomics pipeline identifies small variants, copy-number alterations, and structural variants from sequencing data. The transcriptomics pipeline quantifies gene expression and detects gene fusions from RNA-seq data. The proteomics pipeline identifies and quantifies proteins and post-translational modifications from mass-spectrometry data. The single-cell transcriptomics pipeline characterizes gene expression at cellular resolution. The spatial transcriptomics pipeline characterizes gene expression within its tissue context. The methylomics pipeline profiles genome-wide DNA methylation from bisulfite-sequencing data. Additional pipelines support T-cell receptor profiling, metabolomics, HLA typing, homologous recombination deficiency assessment [19, 20], and neoantigen prediction.

The PromptBio-developed pipelines were evaluated through internal benchmarking and comparison with established external implementations; headline results are summarized in Section 4.3, with full protocols and metrics in Supplementary Note 4.

### 3.2 Analytical Skills

OmicsGenie also provides analytical skills for analyzing processed omics data or user-supplied feature matrices. Differential analysis identifies differentially abundant molecular markers associated with experimental conditions, phenotypes, or other variables of interest. Functional enrichment analysis tests whether selected markers are overrepresented in curated biological pathways, processes, and gene sets. Multi-group comparison identifies differential markers and evaluates expression differences and trends across multiple cohorts or experimental groups. Single-cell analysis supports cell clustering, annotation, marker identification, differential expression analysis, and visualization.

OmicsGenie exposes these skills through defined inputs and outputs, allowing PromptGenie to combine them with upstream processing and downstream interpretation.

### 3.3 Workflow and Pipeline Integration

PromptBio supports complementary mechanisms for integrating external workflows, enabling researchers to extend the platform while reusing trusted community and organizational resources. It dynamically integrates available pipelines from nf-core, a community-driven initiative that develops curated, standardized bioinformatics pipelines [21]. These pipelines can be executed independently or incorporated into multi-step research workflows, expanding PromptBio’s analytical coverage while promoting consistency, reproducibility, and adherence to community best practices.

BYOP enables organizations to register existing scripts or pipelines as callable tools within PromptBio. PromptBio interprets each pipeline’s inputs, outputs, dependencies, and execution requirements, allowing PromptGenie to configure and invoke it either independently or as part of a multi-step research workflow. Organizations can thereby reuse established internal methods alongside PromptBio’s built-in skills and managed execution environments. User-provided pipelines remain distinct from PromptBio-validated skills, and access is restricted to the uploading organization and explicitly authorized parties.

### 3.4 Machine Learning Skills

MLGenie provides the underlying implementation for PromptBio’s machine learning skills, supporting exploratory data analysis as well as classification, regression, and survival analysis. These skills operate on matrix-based data, including high-dimensional omics features, and follow a workflow of preprocessing, feature selection, model development, and multi-omics integration. They can be incorporated into PromptGenie workflows for a broad range of biomedical applications, including biomarker discovery, disease classification, outcome and treatment-response prediction, risk assessment, and investigation of relationships between molecular features and clinical phenotypes. MLGenie integrates established methods for feature selection and predictive modeling. Feature selection comprises filter methods such as univariate filtering, wrapper methods such as backward elimination, and embedded methods including LASSO and random-forest feature importance, with ensemble aggregation and bootstrap sampling used to further improve the stability and reproducibility of selected features. Based on the benchmarking described in Section 4.3, Nullstrap [22] is used as the default feature-selection method. Predictive modeling spans linear models, support vector machines, tree-based methods, and neural networks. Cross-validation and hyperparameter optimization support systematic model comparison and selection, while configurable settings and parallel execution enable reproducible experimentation. The resulting predictive features can subsequently be analyzed using PromptBio’s analytical and knowledge skills to assess their biological functions and supporting literature evidence.

### 3.5 Biomedical Foundation Model Skills

PromptBio integrates selected biomedical foundation models through NVIDIA NIM [23] for structural biology and drug discovery applications. These models enable tasks such as predicting the structure of proteins, RNA, DNA, small molecular ligands, and biomolecular complexes, as well as protein– ligand binding-affinity. These models are exposed through standardized tool interfaces and can be invoked directly by users or selected by PromptGenie as part of multi-step research workflows based on the scientific objective.

### 3.6 Knowledge and Biomarker Skills

MarkerGenie provides the underlying implementation for literature-based biomarker retrieval and interpretation skills. Originally developed to extract disease-specific relationships among biomedical entities from unstructured biomedical literature [24], it identifies relationships involving genes, chemicals such as metabolites and drugs, and microbiome entities. Using MarkerGenie, PromptGenie can retrieve disease-associated entities from the literature and prioritize them by frequency of occurrence, helping researchers rapidly identify commonly reported biomarkers and relationships.

PromptBio further provides knowledge skills that query structured biomedical resources spanning clinical, chemical, pharmacological, genomic, pathway, and target–disease evidence. Representative examples include ClinicalTrials.gov, openFDA, and DailyMed for trial and regulatory records; PubChem, ChEMBL, HMDB, and BindingDB for chemical and bioactivity data; DrugBank, GtoPdb, DGIdb, and PharmGKB for pharmacological and pharmacogenomic annotation; ClinVar, dbSNP, and gnomAD for variant evidence; KEGG and Reactome for pathway context; and Open Targets and Monarch for target–disease associations. Together with MarkerGenie, these skills extend retrieval beyond publications and allow findings to be examined in molecular, pharmacological, genomic, and clinical context.

PromptGenie can integrate literature-derived and structured knowledge with findings from differential analysis, machine learning-based feature selection, or other computational workflows. This integration helps researchers interpret candidate biomarkers, assess the consistency and breadth of supporting evidence, identify potential mechanisms and therapeutic relevance, and prioritize candidates for further computational or experimental investigation. The resulting skills support contextual interpretation of candidate biomarkers while preserving links to the underlying sources and evidence. This evidence-grounded approach makes the resulting interpretations more transparent and traceable than relying on unsupported LLM generation alone.

## 4 Evaluation

We designed the evaluation to assess whether PromptBio can convert biomedical research objectives into accurate, reproducible, and scientifically interpretable outputs. The evaluation was conducted at two complementary levels: (i) system-level performance, measured by PromptGenie’s ability to complete end-to-end bioinformatics analysis and deep-research tasks, and (ii) method-level validity, assessed through key biomedical skills that PromptGenie invokes during execution. Together, these evaluations examine agentic planning and synthesis as well as the trustworthiness of the underlying omics and machine-learning modules. Detailed protocols are provided in Supplementary Notes 2–5.

### 4.1 End-to-End Bioinformatics Analysis

We previously released PromptBio-Bench, a dataset for evaluating whether general-purpose bioinformatics agents can translate natural-language requests and input data into valid analytical outputs [16]. It comprises 244 expert-curated task capsules spanning bioinformatics (131 tasks) and data science (113 tasks), including sequence processing, multi-omics analysis, statistics, machine learning, and visualization. Each capsule contains a natural-language task, domain-appropriate input files, and expert-reviewed reference outputs. The tasks are stratified into low (60), medium (139), and high (45) difficulty tiers. Detailed task design and the evaluation framework are described in the PromptBio-Bench paper; the scoring workflow and agent configurations used here are provided in Supplementary Note 2.

PromptGenie, Biomni, and STELLA received identical tasks. To focus the evaluation on customized analysis, tasks were completed through agent-generated scripts run in the same execution environment; most built-in tools were disabled for Biomni and PromptGenie. Candidate outputs underwent file validation followed by format-specific comparison with expert references; figures and free-text outputs were assessed for scientific-content equivalence using an LLM judge with expert review. A task was counted as completed when the run finished successfully and produced parseable required outputs. For completed tasks, per-file similarity scores were averaged and thresholded at 0.5 to assign binary accuracy; incomplete tasks were assigned an accuracy of 0. Completion rate, output accuracy, and wall-clock latency were recorded over all tasks.

As shown in Figure 3, PromptGenie completed 98.8% of the tasks and achieved a mean accuracy of 0.807 of all completed tasks. Under the tested configurations, its completion rate was comparable to Biomni’s, while its output accuracy was higher than both comparator systems. PromptGenie maintained a high completion rate across all difficulty tiers, whereas output accuracy declined with increased task difficulty for all three systems. The task-level similarity distributions showed the same pattern: high-similarity outputs predominated on low-difficulty tasks but became less frequent with increasing difficulty (Figure S1). Wall-clock latency also increased with task difficulty, although PromptGenie was the fastest system across the reported tiers. Thus, within this evaluation, PromptGenie consistently produced valid outputs across a broad set of tasks, while the decline in output agreement on harder tasks underscores that successful execution does not necessarily ensure analytical correctness. Because this evaluation compares complete agent configurations rather than isolating individual components, the relative results should be interpreted as system-level observations under the reported experimental settings.

**Figure 3:**
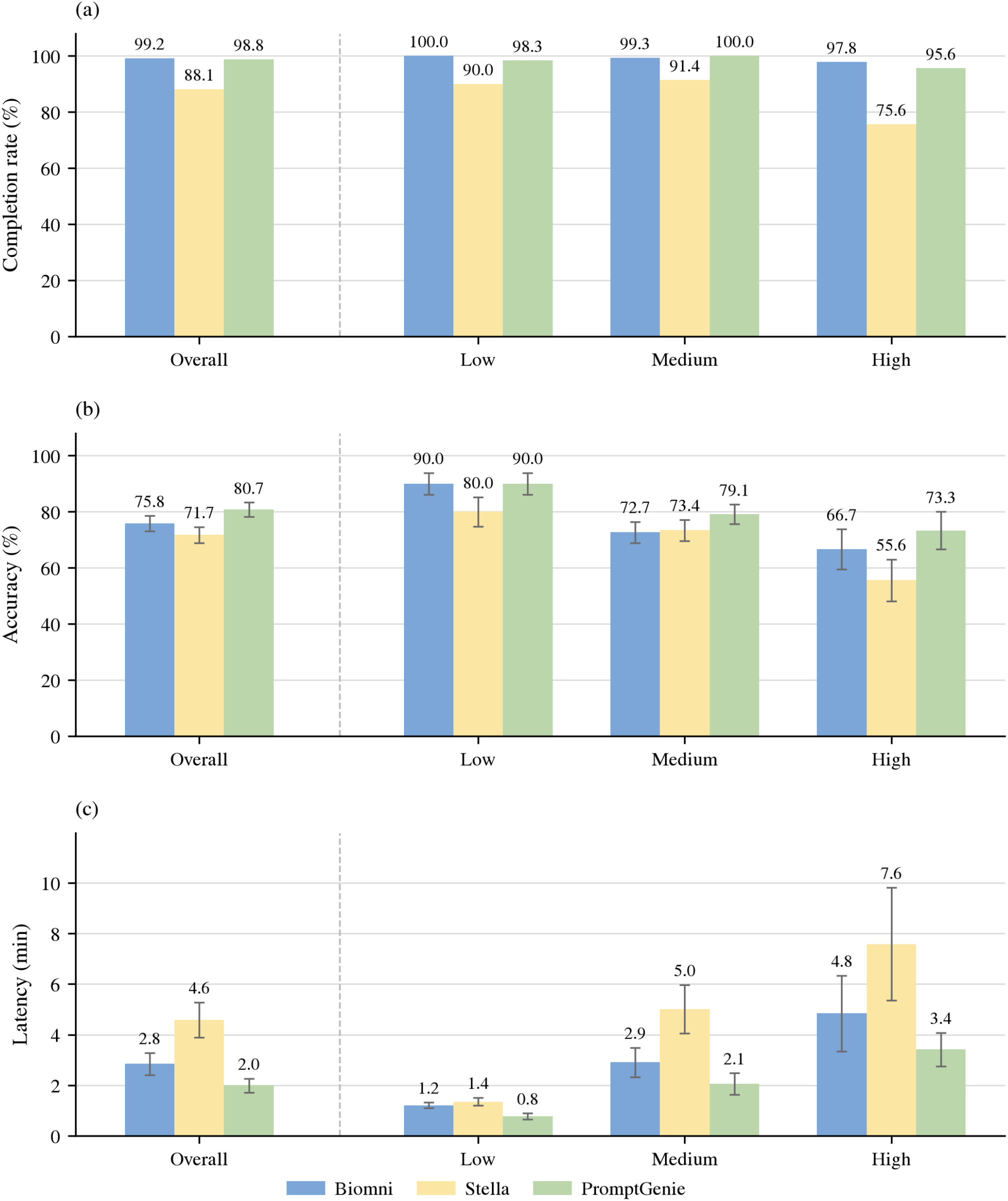
PromptBio-Bench evaluation results for PromptGenie, Biomni, and STELLA. (a) Task completion rate overall and by difficulty. (b) Mean output accuracy with SEM. (c) Mean wall-clock latency with SEM. Bars show means across tasks in each difficulty tier; Overall aggregates all tasks.

### 4.2 Deep-Research Performance

PromptGenie’s evidence-retrieval and synthesis capability was evaluated on the Health subset of DeepResearch Bench II [17], a benchmark derived from expert-written investigative reports. The subset contains 12 tasks—six in English and six in Chinese—evaluated against 782 fine-grained binary rubrics. The rubrics measure information recall, analysis, and presentation, allowing report quality to be assessed at the level of individual factual, inferential, and organizational requirements rather than through a single holistic judgment. An additional blocked component penalizes reliance on withheld source material. PromptGenie was compared with Biomni and Noah.bio using the same task prompts and rubric-based evaluation procedure. Details of agent configurations, latency recording, and the evaluation script are summarized in Supplementary Note 3.

As shown in Figure 4, PromptGenie achieved the highest overall rubric score (0.472) among the systems evaluated. Its relative advantage was concentrated in information recall and analysis, while presentation scores were comparable across systems, suggesting that the difference primarily reflected evidence coverage and analytical content rather than report formatting. PromptGenie’s latency fell between those of Biomni and Noah.bio. Nevertheless, substantial portions of the expert rubrics remained unmet by every system, indicating the continued need for human review. The findings support the use of PromptGenie to produce rapid, structured drafts for evidence-grounded biomedical research, rather than as a substitute for expert synthesis and verification.

**Figure 4:**
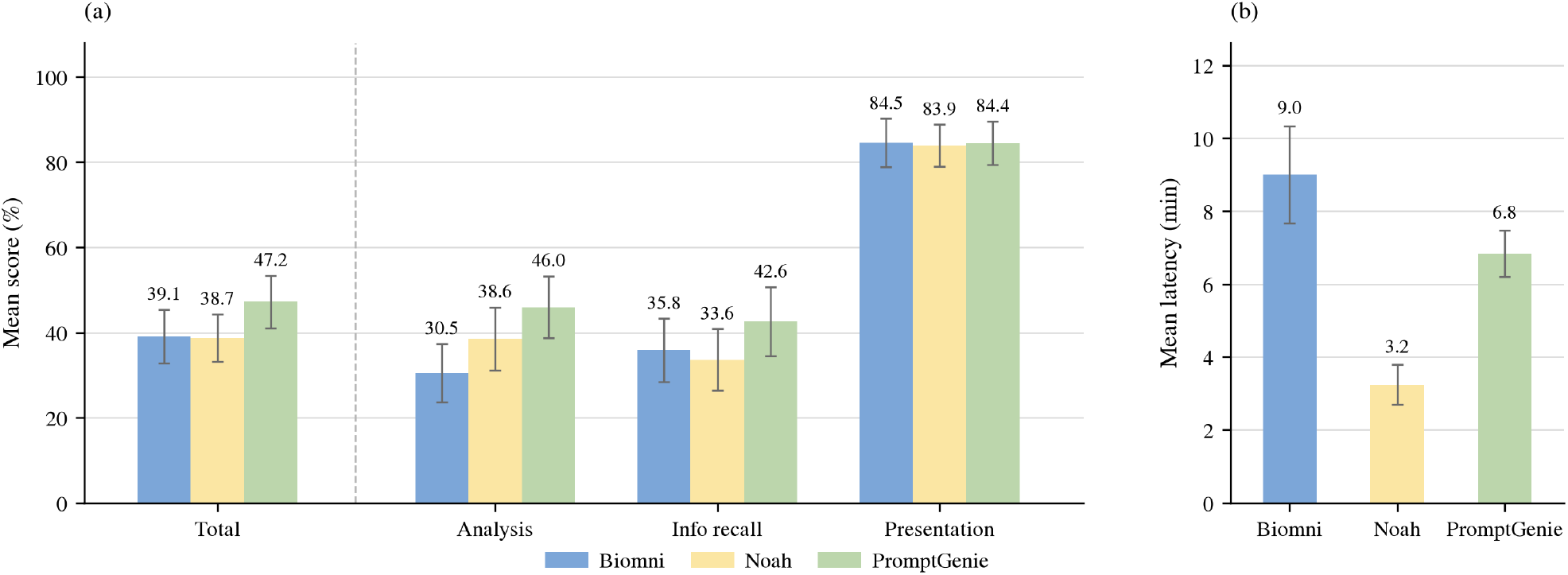
DeepResearch Bench II Health-subset results. (a) Mean rubric scores for the overall (Total) metric and for analysis, information recall, and presentation. All three systems scored 0 on blocked (best), so that component is omitted from the plot. (b) Mean per-task wall-clock latency. Bars show means across the 12 tasks; error bars indicate SEM.

### 4.3 Validation of Biomedical Skills

Because OmicsGenie skills support frequently performed, foundational bioinformatics analyses that require accuracy, reproducibility, and methodological fidelity, we separately validated representative omics-processing and machine-learning modules against established methods and reference implementations. We summarize the principal findings here; detailed datasets, protocols, and performance metrics are provided in Supplementary Notes 4 and 5.

#### Omics processing skills

The genomics, transcriptomics, and proteomics pipelines were evaluated using community benchmarks and established tools (Supplementary Note 4). For short-variant calling, we compared the PromptBio CPU and GPU pipelines with nf-core/sarek using GIAB germline and SEQC2 somatic whole-exome and whole-genome datasets. The pipelines achieved closely comparable accuracy in germline SNP and indel calling and somatic SNV detection. For transcriptomics, we assessed gene quantification and differential expression using SEQC reference RNA-seq datasets with matched RT-qPCR measurements. The results demonstrated high technical reproducibility and substantial concordance with RT-qPCR for gene-level quantification and fold-change estimation. For proteomics, we compared protein identification on the yeast–UPS1 standard LC-MS/MS dataset with results from MaxQuant, Skyline, and MFPaQ, and evaluated quantification against published profiles from a proteogenomic study of chemo-refractory high-grade serous ovarian cancer [25]. The analyses identified a large shared set of proteins and showed strong sample-level correlations with the published protein-expression measurements.

#### MLGenie feature selection

To select the default feature-selection method for MLGenie, we compared two recently developed methods for high-dimensional biological data, Nullstrap [22] and Stabl [26], with two widely used baselines—mutual-information filtering and random-forest importance. We evaluated the four methods on simulated high-dimensional regression tasks spanning a range of feature dimensionalities and noise levels (Supplementary Note 5; Figure S4). Nullstrap achieved the highest AUPRC and F1 score for recovering true signal features. It recovered informative features nearly perfectly under low-noise, lower-dimensional conditions and retained a clear advantage as dimensionality or noise increased. Nullstrap was also the fastest of the four methods (Table S6). These findings motivated its adoption as the default feature-selection method in MLGenie.

Collectively, these results show that PromptGenie’s core biomedical skills align with established analytical practices and can be composed into multi-step research workflows. As with conventional bioinformatics analyses, however, their outputs still require appropriate expert review and interpretation.

## 5 HAQER Regulatory Genomics Case Study

As part of a collaboration investigating the regulatory function of Human Ancestor Quickly Evolved Regions (HAQERs) during human cortical development [18], the PromptBio platform was used to extend the published analysis of human-specific epigenetic features associated with these rapidly evolved sequences. PromptGenie received genomic intervals representing the 1,581 HAQERs, human prefrontal cortex (PFC) tissue-specific and NeuN+ neuronal H3K4me3 peaks, and clusters of human-specific CpG gains (“CpG beacons”), together with a natural-language request to test enrichment, resolve intersections, and connect overlapping regions to genes. The agent inspected the interval data, proposed a researcher-reviewed analysis plan, and generated and executed the complete workflow (Figure 5a). The request, proposed design, and researcher confirmation are provided in Supplementary Note 6.

**Figure 5:**
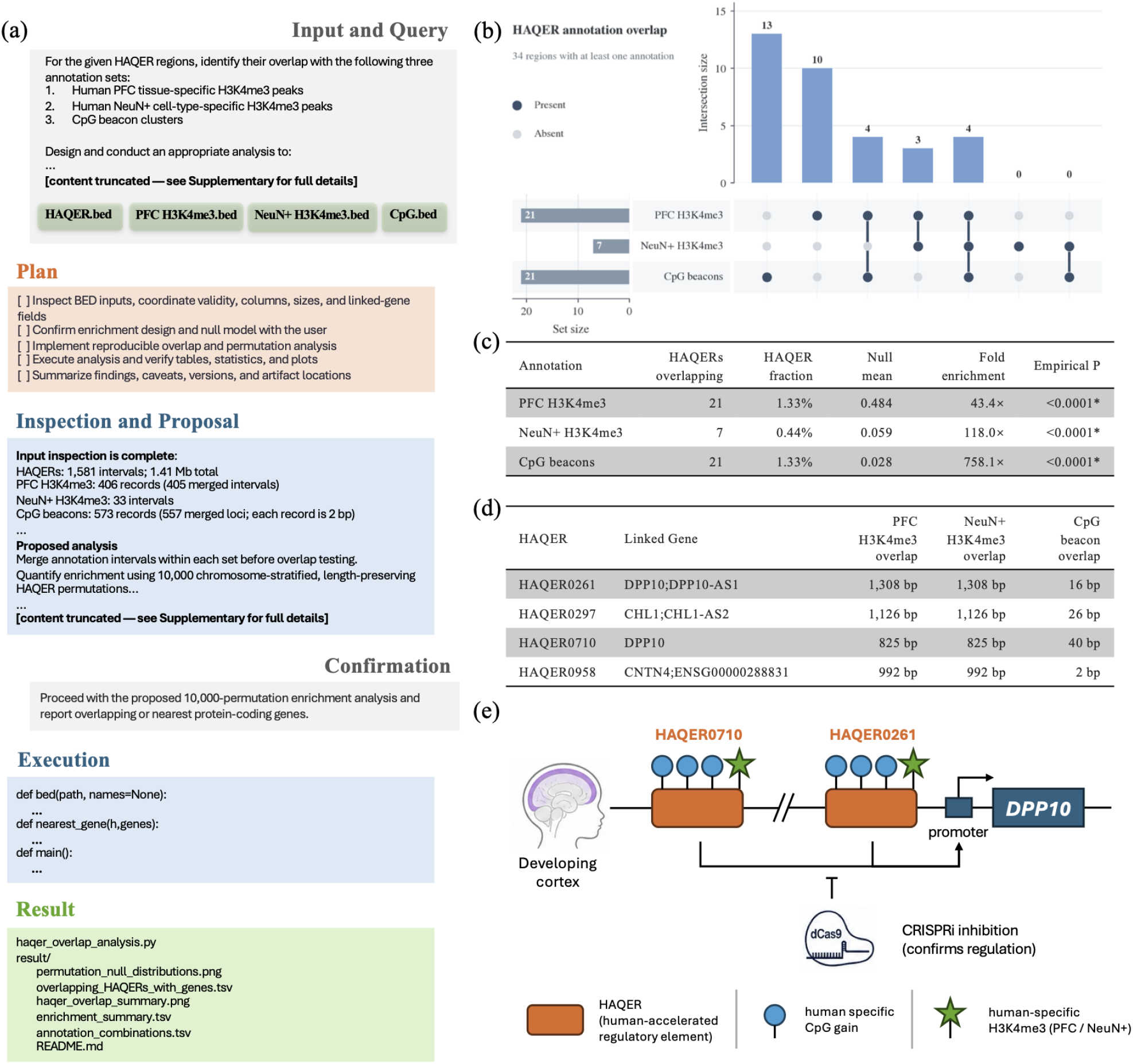
PromptBio-assisted analysis of HAQER regulatory annotations. (a) Starting from a natural-language request and four BED inputs, PromptGenie inspected the data, proposed an enrichment design for researcher confirmation, executed the analysis, and returned reproducible code, tables, and visualizations. (b) UpSet plot summarizing HAQER intersections with PFC-specific H3K4me3, NeuN+-specific H3K4me3, and CpG beacons. (c) Observed overlaps, null expectations, fold enrichments, and empirical permutation *P* values. (d) Representative HAQERs carrying all three annotations and their linked genes. (e) Regulatory context of the CpG-beacon HAQERs HAQER0710 and HAQER0261 at the *DPP10* locus; gene regulation shown here is supported by the collaborative CRISPRi experiments [18].

Chromosome-stratified circular permutation tests showed strong enrichment of HAQERs for all three annotations (Figure 5b,c). A matched random-placement sensitivity analysis produced comparable results, indicating that the enrichment was not attributable to chromosome allocation, HAQER length, or spatial clustering. The 34 annotated HAQERs also exhibited a nested pattern: every NeuN+ overlap occurred within the PFC-overlapping set, and more than half of these neuronal regions also contained CpG beacons. This co-occurrence links rapid human-sequence evolution and CpG gain with chromatin states associated with cortical and neuronal regulation.

Four HAQERs carried all three annotations, including HAQER0297 at *CHL1* and HAQER0261 and HAQER0710 at *DPP10* (Figure 5d). These loci are especially informative because the collaborative study independently showed by CRISPR interference that HAQER0297 and HAQER0202 regulate transcripts at the *CHL1* locus, whereas HAQER0261 and HAQER0710 regulate transcripts at the *DPP10* locus [18]; the *DPP10* example is illustrated in Figure 5e. All four experimentally tested HAQERs are CpG beacons, and three also overlap human-specific neuronal H3K4me3 marks, connecting the computational enrichment to experimentally supported cis-regulatory activity at genes implicated in neurological phenotypes. For the remaining regions, gene-body overlaps and nearest-transcription-start-site assignments provide candidates rather than validated regulatory relationships.

PromptBio completed this researcher-supervised analysis in hours and produced executable code, statistical summaries, region-level and gene-link tables, and publication-ready visualizations. The case study illustrates how the platform contributed a transparent computational component to an active biological collaboration: it transformed a natural-language question into an inspectable analysis while preserving the distinction between statistical association, proximity-based gene assignment, and experimental evidence.

## 6 Security and Operations

PromptBio incorporates operational controls intended to support reliable execution and responsible handling of biomedical data. These controls span workflow observability, access management, data protection, and auditable records. Because deployment requirements differ across research and organizational settings, the applicable controls are configured according to the sensitivity of the data and the policies of the deploying institution.

### 6.1 Operational Monitoring

Centralized logs record task state, execution events, errors, and resource utilization across agent and computational services. Runtime metrics, including processing time, memory use, and failure rates, support capacity planning, performance analysis, and identification of abnormal behavior. Configurable alerts notify operators of events such as failed tasks, unavailable services, or resource exhaustion, enabling investigation and recovery while preserving the associated execution record.

### 6.2 Data and Service Security

PromptBio applies layered controls to data, services, and computational tasks. Data are encrypted at rest using AES-256 and protected in transit using TLS. Role-based access control limits project data and platform operations to authorized users, while task-scoped permissions restrict the data and credentials available during execution. Service and API access uses OAuth 2.0-based authentication, and organization-specific pipelines and artifacts remain isolated from users who have not been explicitly authorized. Together, these measures reduce unnecessary data exposure across interactive, agent-mediated, and computational workflows.

### 6.3 Compliance Readiness and Auditability

The platform is designed to support deployments subject to institutional governance and frameworks such as HIPAA and SOC 2. Relevant mechanisms include access control, encryption, configurable data-retention practices, and audit records of user activity, data access, workflow execution, and administrative changes. These records can support incident investigation, reproducibility, and organizational compliance assessment. Compliance nevertheless depends on the complete deployment—including institutional policies, operating procedures, infrastructure configuration, and contractual controls—and should not be inferred from platform features alone.

## 7 Discussion and Future Directions

PromptBio is centered on an agent harness for computational biomedical research. PromptGenie integrates reasoning and planning, subagent delegation, validated skills, custom code generation, managed execution, and evidence synthesis within a common control loop. This architecture maintains continuity of the research objective across literature and data discovery, analysis, interpretation, and reporting.

A central design consideration is the balance between analytical flexibility and scientific control. Validated skills provide maintained implementations for established tasks, while generated code and imported pipelines extend the platform beyond the existing skill library. Inspectable plans, managed environments, execution records, and provenance make each path reviewable. Human oversight remains integral to the design: researchers can refine plans, review methods and intermediate outputs, and intervene when evidence is ambiguous or an analytical decision could affect the scientific conclusion.

The evaluations examine complementary aspects of the platform. On PromptBio-Bench, PromptGenie completed nearly all tasks and achieved higher output accuracy and lower latency than the comparison agents. On the Health subset of DeepResearch Bench II, its relative advantage was concentrated in information recall and analysis rather than presentation. Representative omics and machine-learning skills produced results consistent with established methods, supporting their use as building blocks within agentic workflows. The HAQER case study further demonstrates how planning, custom analysis, managed execution, and researcher review can be coordinated in a hypothesis-driven investigation.

These evaluations nevertheless have limitations. They cover only a subset of computational biomedical research; reported performance depends on the underlying language models, which were not matched across systems; single-pass scores do not capture run-to-run variability; and the benchmarks assess final artifacts rather than intermediate reasoning or generated code. Cost and latency were also not systematically compared and remain practical constraints. Broader benchmarks are needed across diverse questions, data types, and analytical settings.

Future development will focus on the following areas:

### Expanded Scientific Skills and Models

The skill library will add support for additional modalities and methods, including ATAC-seq, ChIP-seq, and Hi-C, and broaden the analyses available for existing modalities such as spatial transcriptomics. Additional biomedical foundation models and GPU-accelerated inference services will extend support for structural biology, drug discovery, and related applications.

### Integrated Data and Knowledge Resources

Expanding literature, dataset, and biomedical knowledge resources will allow PromptGenie to connect project-specific results with relevant molecular, pharmacological, genomic, and clinical evidence within the same research trajectory.

### Experience-Informed Skill Development

Successful custom analyses, researcher-corrected plans, and execution recoveries can identify recurring methods that warrant conversion into reusable skills. Promotion will remain subject to expert review and validation, allowing the platform to learn from use without treating prior execution alone as evidence of scientific validity.

### Scalable and Reusable Research Workflows

We will improve resource-aware scheduling across CPU and GPU environments, recovery of long-running tasks, and interoperability with external tools and workflow systems. Interactive result exploration and visual workflow controls will support inspection, refinement, and reuse of completed analyses.

### Component and End-to-End Evaluation

New benchmarks will assess planning, retrieval, skill selection, execution recovery, and synthesis both individually and across complete research trajectories. Evaluation with external datasets, independent users, and domain-expert review will test robustness and generalizability.

Together, these developments will extend the scientific reach of the harness while preserving the transparency, reproducibility, and human accountability required for biomedical research.

## Acknowledgments

We thank our external trial users for their valuable feedback, which directly contributed to improving the platform. We are grateful to Yashodara Abeykoon and Prof. Alex A. Pollen (University of California, San Francisco) for providing materials that enabled the HAQER regulatory-genomics case study.

