## Supplementary Information for "PromptBio: An Agentic Platform for End-to-End Computational Biomedical Research"

#### Supplementary Note 1. DataGenie Supported Data Sources

DataGenie discovers and acquires public biomedical datasets for PromptGenie workflows. Table S1 summarizes the fifteen commonly used repositories currently incorporated, spanning sequencing archives (SRA, GEO, ArrayExpress, and GTEx), proteomics resources (PRIDE, iProX, SCP, PDC, CPTAC), metabolomics (MetaboLights), cancer genomics collections (GDC, cBioPortal, TCGA), and pharmacogenomic and functional-genomics resources (GDSC and DepMap). Dataset and sample counts reflect the indexed inventory available to DataGenie at the time of this writing. For PRIDE, iProX, SCP, and GDSC, the number of samples cannot be directly derived from file counts and is therefore marked as NA.

Table S1: Database summary in DataGenie. Commonly used databases are listed with dataset and sample counts. For PRIDE, iProX, SCP, and GDSC, the number of samples cannot be directly derived from file counts and is marked as NA.

| Database | Number of datasets | Number of samples |
|---|---:|---:|
| SRA | 636,336 | 37,902,411 |
| GEO | 306,361 | 10,029,625 |
| ArrayExpress | 80,215 | 2,959,988 |
| PRIDE | 37,857 | NA |
| iProX | 6,452 | NA |
| MetaboLights | 2,838 | 428,399 |
| GDC | 1,790 | 1,200,391 |
| cBioPortal | 1,376 | 578,469 |
| TCGA (raw) | 1,015 | 445,343 |
| SCP | 984 | NA |
| PDC | 214 | 16,750 |
| CPTAC (raw) | 197 | 25,321 |
| TCGA (processed) | 169 | 56,261 |
| DepMap | 160 | 1,067 |
| GTEx | 67 | 19,374 |
| CPTAC (processed) | 57 | 7,482 |
| GDSC | 32 | NA |

### Supplementary Note 2. PromptBio-Bench Evaluation Protocol

Task design, difficulty stratification, input/reference packaging, and the full evaluation framework are described in the PromptBio-Bench paper [1]. Here we summarize the scoring workflow and agent configurations used for the comparisons reported in the main text.

**Scoring workflow.** A task was flagged as completed if the agent run finished successfully, produced the required output artifacts, and those outputs could be parsed by the evaluation pipeline. For each completed task, a similarity score was computed by comparing each generated file with the corresponding expert reference: structured files used format-specific comparison, and unstructured outputs such as figures and free-text summaries were scored with an LLM judge (GPT-5.4). Further details of these comparison procedures are provided in the PromptBio-Bench paper [1]. When a task produced multiple files, per-file similarity scores were averaged to obtain a single task-level similarity. Binary accuracy was then assigned by thresholding the task-level similarity at 0.5 (similarity $\geq 0.5$ scored as 1; otherwise 0). Incomplete tasks were assigned a similarity of 0 and therefore an accuracy of 0. Completion rate, mean accuracy, and wall-clock latency were reported over all tasks, overall and by difficulty tier.

**Comparison agents.** We evaluated three general-purpose bioinformatics agents: Biomni (v0.0.8), STELLA (v1.0.0), and PromptGenie (v0.7.0). To focus on customized analysis and improve comparability, tasks were completed through agent-generated scripts run in the same execution environment, with most built-in tools disabled for Biomni and PromptGenie. Biomni used Claude Sonnet 4 as its LLM backend. STELLA used the hybrid model strategy described in its paper: the Dev Agent and Tool Creation Agent used Claude Sonnet 4, while the Manager Agent and Critic Agent used Gemini 2.5 Pro. PromptGenie used GPT-5.6 Sol as its LLM backend. Thus, differences may reflect both agent architecture and LLM backend. Comparisons under more closely matched model conditions are reported in the original PromptBio-Bench study [1].

**Similarity-score distributions.** Figure S1 summarizes the distribution of task-level similarity scores in four bins—$[0, 0.25)$, $[0.25, 0.5)$, $[0.5, 0.75)$, and $[0.75, 1]$—stratified by difficulty. On the low-difficulty tier, scores concentrate in the high-similarity bin ($[0.75, 1]$), with little mass in the intermediate ranges. As difficulty increases, the high-similarity share declines and mass shifts into the lower and middle bins for all three agents; PromptGenie retains the largest high-similarity share on the medium- and high-difficulty tiers.

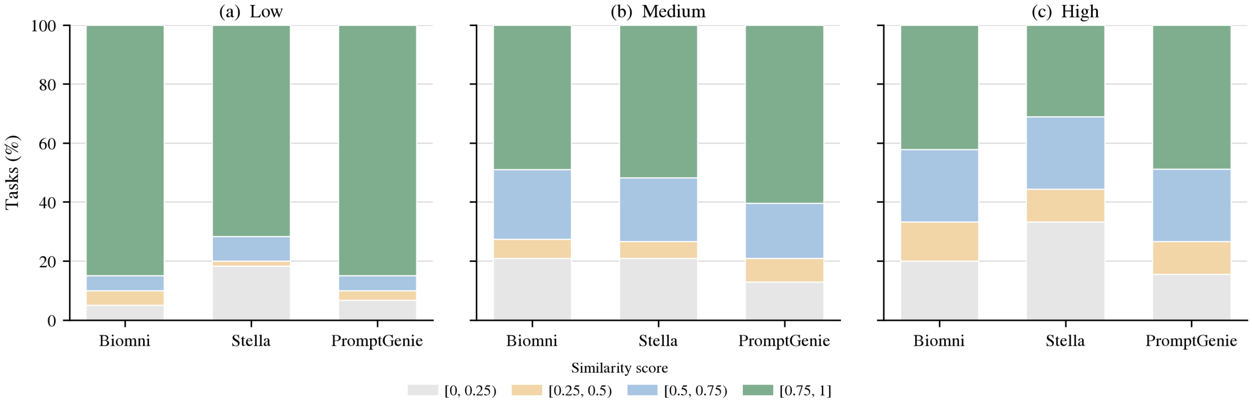


Figure S1: Distribution of PromptBio-Bench task-level similarity scores for Biomni, STELLA, and PromptGenie, stratified by difficulty. Bars show the percentage of tasks whose averaged per-file similarity fell in each bin: $[0, 0.25)$, $[0.25, 0.5)$, $[0.5, 0.75)$, or $[0.75, 1]$. Panels: (a) low, (b) medium, and (c) high difficulty.

### Supplementary Note 3. Deep-Research Evaluation Protocol

Benchmark design, task construction, and the rubric inventory are described in the DeepResearch Bench II paper [2]. Here we summarize the subset, agent configurations, latency recording, and scoring procedure used for the comparisons reported in the main text.

**Benchmark subset.** We evaluated systems on the Health subset of DeepResearch Bench II, comprising 12 tasks (six in English and six in Chinese) and 782 fine-grained binary rubrics. The rubrics score information recall, analysis, and presentation; an additional blocked component penalizes reliance on withheld source material. All systems received the same task prompts.

**Comparison agents.** PromptGenie (v0.7.0) was run on the PromptBio platform. Biomni and Noah.bio were accessed through their public websites in August 2026 and evaluated with each site's default model settings. Wall-clock time from prompt submission to completed report was recorded for every task.

**Scoring.** Generated reports were scored with the official DeepResearch Bench II evaluation script (`https://github.com/imlrz/DeepResearch-Bench-II`). Per-task rubric scores were aggregated into the Total metric and the analysis, information-recall, and presentation components reported in the main text.

### Supplementary Note 4. Omics Processing Skill Validation

We validated representative PromptBio omics skills against community benchmarks and established tools. Genomics short-variant calling, transcriptomics quantification, and proteomics identification/quantification are reported below.

#### Genomics: Germline and Somatic Short-Variant Calling

We evaluated WES and WGS short-variant calling in germline and somatic settings by comparing three pipeline configurations under matched inputs and callers: nf-core/sarek v3.5.1 [3], PromptBio_CPU, and PromptBio_GPU (NVIDIA Parabricks-accelerated). Germline calling used GATK HaplotypeCaller; somatic calling used GATK Mutect2. Only `PASS` variants were scored. SNVs/SNPs and indels were evaluated separately with Illumina `hap.py` (germline) or `som.py` (somatic), reporting recall, precision, and F1 within high-confidence regions.

**Germline benchmark (GIAB HG001/NA12878, GRCh38).** Truth calls and high-confidence intervals were taken from GIAB HG001 v4.2.1 [4]. WES evaluation used two independent Garvan/Illumina exome libraries (NIST7035 and NIST7086; HiSeq2500, 101 bp paired-end, ∼100×), restricted to the intersection of GIAB high-confidence regions and the exome capture targets. WGS evaluation used NA12878 (SRR6794144; HiSeq4000, 150 bp paired-end, ∼50×) over the full high-confidence BED.

**Somatic benchmark (SEQC2 HCC1395/HCC1395BL, GRCh38).** Truth calls and high-confidence intervals were taken from the SEQC2 Somatic Mutation Working Group release [5]. WES evaluation used five independent tumor/normal exome pairs from SRA study SRP162370 spanning a range of depths (∼111–516× tumor), restricted to SEQC2 high-confidence regions intersected with the SEQC2 exome target BED. WGS evaluation used one tumor/normal pair (WGS_NC_T_1 / WGS_NC_N_1) over the full high-confidence regions.

Germline performance was highly concordant across sarek, PromptBio_CPU, and PromptBio_GPU in both WES and WGS (Table S2). WES SNP F1 scores clustered between 0.9369 and 0.9431 across the two libraries; WES indel F1 scores were lower, as expected (0.7149–0.7238), with only marginal pipeline differences. WGS accuracy was substantially higher than WES for both

SNPs (F1 > 0.976) and indels (F1 > 0.917), with PromptBio_GPU showing the strongest overall WGS SNP and indel F1.

Table S2: Germline short-variant calling on GIAB HG001/NA12878. Metrics from `hap.py` within high-confidence (and WES target-restricted) regions.

| Sample | Pipeline | Recall | Precision | F1 |
|---|---|---|---|---|
| *WES SNPs* | | | | |
| NIST7035 | sarek_3.5.1 | 0.9083 | 0.9674 | 0.9369 |
| NIST7035 | PromptBio_CPU | 0.9096 | 0.9696 | 0.9386 |
| NIST7035 | PromptBio_GPU | 0.9095 | 0.9706 | 0.9391 |
| NIST7086 | sarek_3.5.1 | 0.9162 | 0.9681 | 0.9414 |
| NIST7086 | PromptBio_CPU | 0.9175 | 0.9698 | 0.9429 |
| NIST7086 | PromptBio_GPU | 0.9175 | 0.9702 | 0.9431 |
| *WES indels* | | | | |
| NIST7035 | sarek_3.5.1 | 0.7411 | 0.6913 | 0.7153 |
| NIST7035 | PromptBio_CPU | 0.7407 | 0.6908 | 0.7149 |
| NIST7035 | PromptBio_GPU | 0.7430 | 0.6889 | 0.7149 |
| NIST7086 | sarek_3.5.1 | 0.7555 | 0.6947 | 0.7238 |
| NIST7086 | PromptBio_CPU | 0.7549 | 0.6939 | 0.7231 |
| NIST7086 | PromptBio_GPU | 0.7567 | 0.6935 | 0.7237 |
| *WGS SNPs* | | | | |
| NA12878 | sarek_3.5.1 | 0.9862 | 0.9662 | 0.9761 |
| NA12878 | PromptBio_CPU | 0.9862 | 0.9672 | 0.9766 |
| NA12878 | PromptBio_GPU | 0.9873 | 0.9845 | 0.9859 |
| *WGS indels* | | | | |
| NA12878 | sarek_3.5.1 | 0.9133 | 0.9216 | 0.9174 |
| NA12878 | PromptBio_CPU | 0.9133 | 0.9222 | 0.9177 |
| NA12878 | PromptBio_GPU | 0.9164 | 0.9545 | 0.9351 |

WES somatic SNV performance was strong across all five SEQC2 pairs (Table S3): PromptBio_CPU achieved the highest F1 in three pairs and PromptBio_GPU in two, while sarek remained competitive throughout. WES somatic indels were not reported separately because too few high-confidence evaluable indels were available in the targeted regions for stable cross-pipeline comparison. On the WGS somatic pair, all three pipelines showed high SNV precision (F1 0.9099–0.9258; highest F1 for PromptBio_CPU) and comparable indel performance (F1 0.7788–0.8036; highest F1 for PromptBio_GPU).

Across GIAB germline and SEQC2 somatic benchmarks, PromptBio_CPU and PromptBio_GPU were comparable to sarek v3.5.1 for germline SNP/indel calling and somatic SNV detection in both WES and WGS, with reliable WGS somatic indel performance. Differences among pipelines were generally small relative to the WES–WGS accuracy gap.

#### Transcriptomics: Gene Quantification and Differential Expression

We validated the PromptBio Transcriptomics Pipeline on two SEQC reference datasets with matched RNA-seq and RT-qPCR profiles: GSE47792 [6] and GSE83402 [7]. Each dataset includes the standardized human reference RNAs MAQCA (Universal Human Reference RNA) and MAQCB (Human Brain Reference RNA), each sequenced in technical duplicate.

RNA-seq reads were processed to gene-level transcripts per million (TPM); analyses were

Table S3: Somatic short-variant calling on SEQC2 HCC1395/HCC1395BL. Metrics from `som.py` within high-confidence (and WES target-restricted) regions. WES somatic indels omitted (insufficient evaluable truth indels).

| Sample | Pipeline | Recall | Precision | F1 |
|---|---|---|---|---|
| *WES SNVs* | | | | |
| WES_NC | sarek_3.5.1 | 0.8481 | 0.9666 | 0.9035 |
| WES_NC | PromptBio_CPU | 0.8602 | 0.9784 | 0.9155 |
| WES_NC | PromptBio_GPU | 0.8775 | 0.9649 | 0.9191 |
| WES_LL | sarek_3.5.1 | 0.7084 | 0.9785 | 0.8218 |
| WES_LL | PromptBio_CPU | 0.7575 | 0.9400 | 0.8390 |
| WES_LL | PromptBio_GPU | 0.7714 | 0.9322 | 0.8442 |
| WES_IL | sarek_3.5.1 | 0.8766 | 0.9864 | 0.9283 |
| WES_IL | PromptBio_CPU | 0.8965 | 0.9738 | 0.9335 |
| WES_IL | PromptBio_GPU | 0.9215 | 0.9247 | 0.9231 |
| WES_EA | sarek_3.5.1 | 0.8844 | 0.9535 | 0.9176 |
| WES_EA | PromptBio_CPU | 0.8991 | 0.9396 | 0.9189 |
| WES_EA | PromptBio_GPU | 0.9146 | 0.8185 | 0.8639 |
| WES_FD | sarek_3.5.1 | 0.7101 | 0.9717 | 0.8205 |
| WES_FD | PromptBio_CPU | 0.7903 | 0.9328 | 0.8557 |
| WES_FD | PromptBio_GPU | 0.8154 | 0.8491 | 0.8319 |
| *WGS SNVs* | | | | |
| WGS_NC | sarek_3.5.1 | 0.8568 | 0.9890 | 0.9182 |
| WGS_NC | PromptBio_CPU | 0.8663 | 0.9941 | 0.9258 |
| WGS_NC | PromptBio_GPU | 0.8361 | 0.9979 | 0.9099 |
| *WGS indels* | | | | |
| WGS_NC | sarek_3.5.1 | 0.8958 | 0.7167 | 0.7963 |
| WGS_NC | PromptBio_CPU | 0.9167 | 0.6769 | 0.7788 |
| WGS_NC | PromptBio_GPU | 0.9375 | 0.7031 | 0.8036 |

restricted to protein-coding genes. TPM values were log-transformed. RT-qPCR genes were retained if Cq was between 11 and 32; RNA-seq genes were required to have TPM $> 0$ in all replicates, with extreme outliers removed. Performance was assessed as squared Pearson correlation ($R^2$) for (i) technical-replicate log-TPM concordance, (ii) RNA-seq versus RT-qPCR log-expression concordance on overlapping genes, and (iii) concordance of MAQCA–MAQCB log fold changes between RNA-seq and RT-qPCR.

Technical reproducibility was high in both datasets ($R^2 = 0.961$–$0.975$ across MAQCA/MAQCB replicates; Table S4). Agreement with RT-qPCR was moderate to strong for absolute log expression ($R^2 = 0.627$–$0.688$; 12,013 overlapping genes in GSE47792 and 13,738 in GSE83402) and stronger for differential expression, with log-fold-change $R^2 = 0.851$ (GSE47792) and 0.859 (GSE83402).

Table S4: Transcriptomics validation on SEQC datasets with matched RT-qPCR. $R^2$ denotes squared Pearson correlation of log-transformed values.

| Dataset | Comparison | $R^2$ (MAQCA) | $R^2$ (MAQCB) | $R^2$ (fold change) |
|---|---|---|---|---|
| GSE47792 | Technical replicates (log TPM) | 0.961 | 0.966 | — |
| GSE47792 | RNA-seq vs RT-qPCR (log expression) | 0.636 | 0.627 | — |
| GSE47792 | RNA-seq vs RT-qPCR (log fold change) | — | — | 0.851 |
| GSE83402 | Technical replicates (log TPM) | 0.975 | 0.964 | — |
| GSE83402 | RNA-seq vs RT-qPCR (log expression) | 0.688 | 0.668 | — |
| GSE83402 | RNA-seq vs RT-qPCR (log fold change) | — | — | 0.859 |

Across two independent SEQC benchmarks, the PromptBio Transcriptomics Pipeline showed excellent technical reproducibility and substantial concordance with RT-qPCR for gene-level quantification and fold-change estimation.

##### Proteomics: Protein Identification and Quantification

Protein identification was benchmarked on a yeast–UPS1 LC-MS/MS standard (PRIDE: PXD001819) [8]. Proteins identified by the PromptBio proteomics pipeline were compared with MaxQuant (Intensity and LFQ), Skyline, and MFPaQ. Across the five methods, 606 proteins were shared by all pipelines; PromptBio identified 801 proteins in total, with set sizes for the reference tools ranging from 790 (Skyline) to 1,091 (MFPaQ). The large core intersection supports consistency of the identification workflow (Figure S2).

Quantification was evaluated on the proteogenomic chemo-refractory high-grade serous ovarian cancer dataset [9]. PromptBio protein expression was compared with publication-reported profiles using Pearson correlation. In the frozen and FFPE validation cohorts, 100% of samples achieved $R \geq 0.8$, and 93.3% of samples in the FFPE discovery cohort met the same threshold (Table S5). Figure S3 shows the correlation matrix for the FFPE validation cohort ($n = 20$) versus PDC reference data; matched sample pairs form a strong diagonal ($R = 0.82$–$0.96$), with low off-diagonal correlations.

#### Supplementary Note 5. MLGenie Feature-Selection Validation

To choose a feature-selection method for PromptBio's machine-learning skills, we required (1) mature Python/R implementations, (2) scalability to high-dimensional data, (3) support for classification, regression, and survival analysis, (4) sparse and interpretable feature scores, (5) limited manual hyperparameter tuning, and (6) an automatic recommendation for the number of selected features.

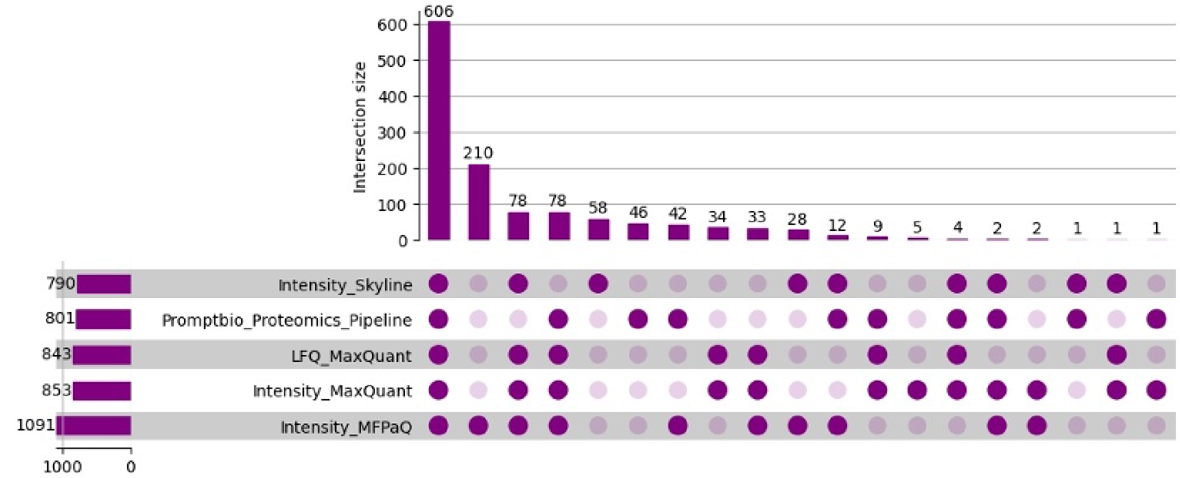


Figure S2: UpSet plot of proteins identified by PromptBio and reference tools (MaxQuant Intensity/LFQ, Skyline, MFPaQ) on the yeast–UPS1 standard (PRIDE: PXD001819). Horizontal bars show set sizes; vertical bars show intersection sizes for each combination of pipelines.

Table S5: Correlation of protein expression levels between PromptBio and the Cell article. Values are the proportion of samples with Pearson correlation coefficient $R$ at or above the indicated threshold.

| Cohort | $R \geq 0.9$ | $R \geq 0.8$ | $R \geq 0.7$ |
|---|---|---|---|
| FFPE Discovery Cohort | 44.3% | 93.3% | 97.6% |
| Frozen Validation Cohort | 51.4% | 100.0% | 100.0% |
| FFPE Validation Cohort | 75.0% | 100.0% | 100.0% |

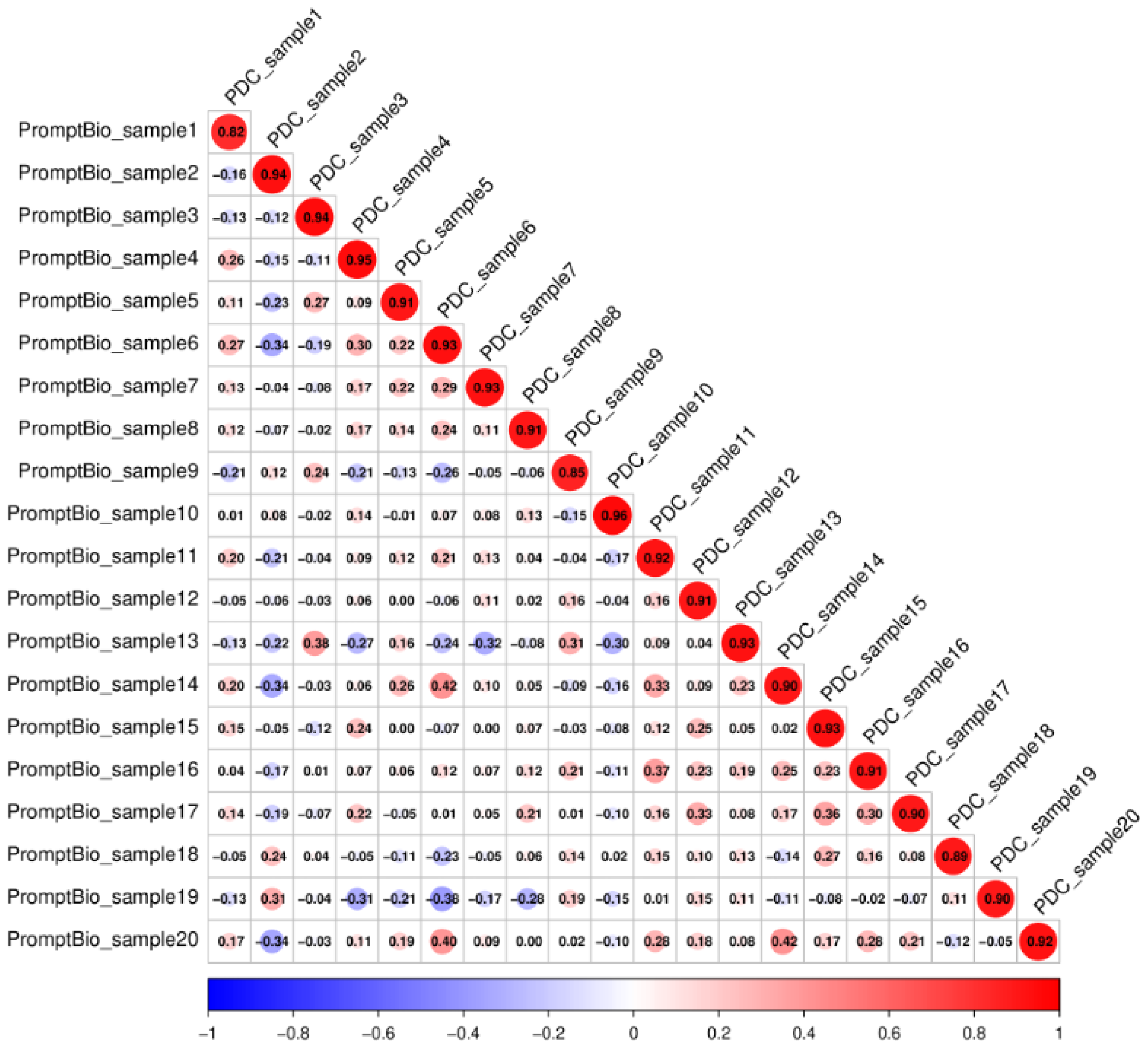

Figure S3: Correlation matrix comparing protein expression levels between PromptBio results and PDC data for the FFPE validation cohort ($n = 20$). Each cell represents a Pearson correlation coefficient between a sample pair, colored by correlation level.

Guided by these criteria, we compared Nullstrap [10] with Stabl [11] and two common baselines—mutual-information filtering and random-forest importance—under controlled simulations. Full simulation designs, additional tasks, and extended results are reported previously [12].

We summarize regression results here. Each simulation used $n = 300$ samples and a feature matrix of dimensionality $p \in \{1000, 3000, 10000\}$, of which $p_s = 10$ features were true signal features associated with the response and the remainder were non-informative. Noise was controlled at three levels (low, medium, and high; $\alpha \in \{0.2, 0.4, 0.6\}$). Mutual information and random forest ranked features by univariate association or model-based importance, respectively, and the top 10 features were retained for comparison with Nullstrap and Stabl. Recovery of the true signal features was assessed by AUPRC and F1 score.

Across scenarios, Nullstrap achieved the highest AUPRC and F1, nearly perfectly recovering informative features at low noise and lower dimensionality, and retaining a clear advantage as $p$ or noise increased (Figure S4). Stabl generally ranked second, followed by random forest, with mutual information weakest throughout. Nullstrap was also the fastest method (mean ∼1–9 s for $p = 1000$–$10000$), whereas Stabl was the slowest and scaled least favorably with dimensionality (Table S6). These results motivated adoption of Nullstrap for feature selection within MLGenie.

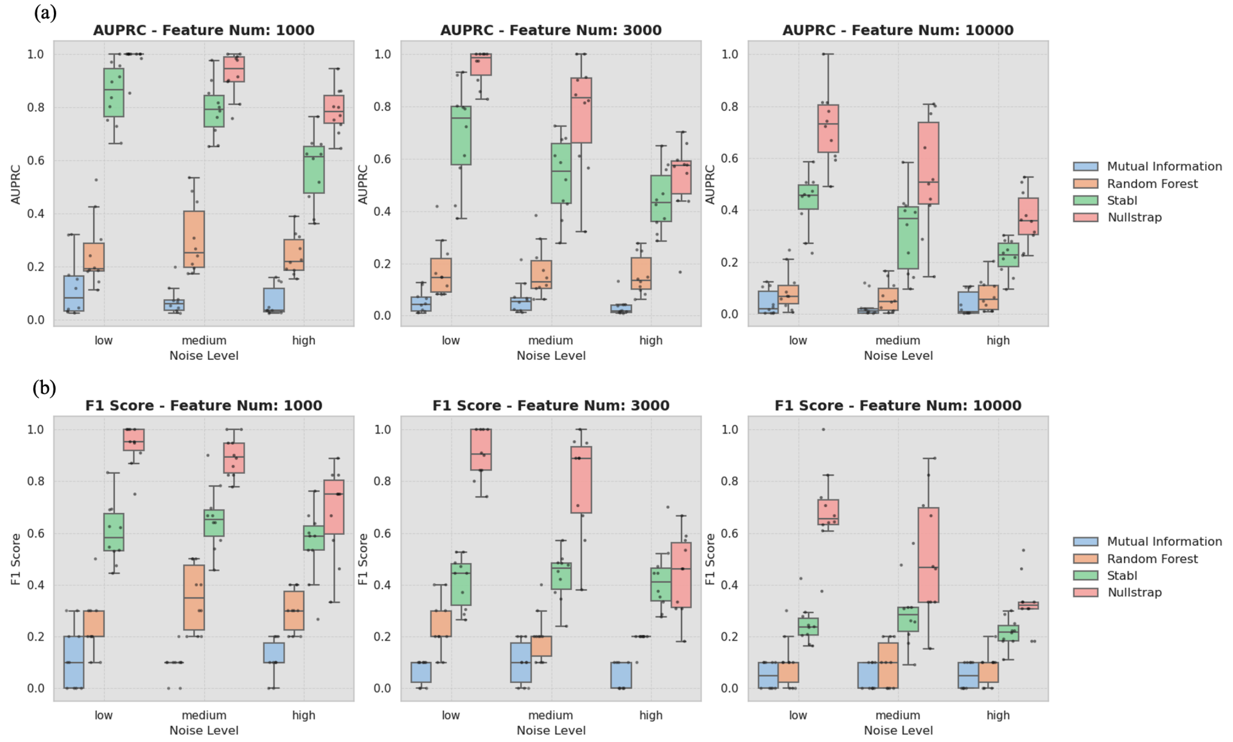


Figure S4: Feature-selection performance on simulated linear regression tasks ($n = 300$; $p_s = 10$ informative features). Panels show (a) AUPRC and (b) F1 score for mutual information, random forest, Stabl, and Nullstrap across feature dimensionalities $p \in \{1000, 3000, 10000\}$ and noise levels (low, medium, high). Boxes summarize 10 simulation replicates; points show individual replicates.

Table S6: Computation time (in seconds) for feature selection methods across simulation settings. Mean wall-clock time (± standard deviation) is reported for each method, averaged over 10 simulation replicates. Results are shown for linear regression tasks with varying feature dimensions ($p = 1000$, 3000, and 10000; $n = 300$).

| Method | $p = 1000$ | $p = 3000$ | $p = 10000$ |
|---|---|---|---|
| Mutual information | $1.96 \pm 0.02$ | $5.87 \pm 0.05$ | $19.52 \pm 0.19$ |
| Random forest | $41.03 \pm 0.77$ | $122.16 \pm 1.90$ | $406.27 \pm 7.78$ |
| Stabl | $92.60 \pm 2.13$ | $348.66 \pm 29.11$ | $1644.52 \pm 151.70$ |
| Nullstrap | $1.17 \pm 0.16$ | $3.08 \pm 0.40$ | $8.50 \pm 0.75$ |

### Supplementary Note 6. HAQER Regulatory-Annotation Analysis

The analysis supporting Figure 5 was initiated from a natural-language request and four BED inputs (HAQERs, PFC-specific H3K4me3, NeuN+-specific H3K4me3, and CpG beacon clusters). PromptGenie inspected the interval files, proposed an enrichment and intersection design, and paused for researcher confirmation before execution (Figure 5a). The request, the agent's first response, and the researcher's confirmation are reproduced verbatim below. Enrichment statistics, annotation intersections, representative gene links, and biological context are reported in Figure 5b–e.

#### Researcher request

```
For the given HAQER regions, identify their overlap with the following three annotation sets:
1. Human PFC tissue-specific H3K4me3 peaks
2. Human NeuN+ cell-type-specific H3K4me3 peaks
3. CpG beacon clusters

Design and conduct an appropriate analysis to:
1. determine whether HAQERs are significantly enriched for each of the three annotation sets.
2. characterize the intersections among the three annotation sets within the HAQER regions.

Formulate and present the results with appropriate summaries, tables, and visualizations that
    clearly illustrate the key findings. For the overlapping HAQER regions, report the
    corresponding linked genes where available.
```

#### Agent inspection and proposed design

```
Input inspection is complete:
HAQERs: 1,581 intervals; 1.41 Mb total
PFC H3K4me3: 406 records (405 merged intervals)
NeuN+ H3K4me3: 33 intervals
CpG beacons: 573 records (557 merged loci; each record is 2 bp)
All coordinates are valid BED intervals on chr-prefixed hg38 chromosomes.
Preliminary overlaps involve 21 HAQERs with PFC, 7 with NeuN+, and 21 with CpG beacons.
The supplied files contain no explicit linked-gene field.

Proposed analysis
Merge annotation intervals within each set before overlap testing.
Quantify enrichment using 10,000 chromosome-stratified, length-preserving HAQER permutations,
    excluding assembly gaps and retaining the observed chromosome distribution.
```

```
Use the number of HAQERs overlapping each annotation as the primary statistic; also report
    covered base pairs and empirical fold enrichment, one-sided permutation P-values, and
    BH-adjusted q-values.
Characterize all seven nonempty combinations of the three annotations per HAQER, with
    UpSet-style and genome-level visualizations.
Link overlapping HAQERs to genes using hg38 gene annotations:
overlapping genes;
otherwise nearest protein-coding gene and TSS distance.
Please confirm this design before I run the full analysis. If you prefer a stricter
    regulatory-gene definition---such as promoter overlap only---select that instead.
```

##### Researcher confirmation

```
Proceed with the proposed 10,000-permutation enrichment analysis and report overlapping or
    nearest protein-coding genes.
```
